# Homing Choices of Breast Cancer Cells Revealed by Tissue Specific Invasion and Extravasation Lab-on-a-chip Platforms

**DOI:** 10.1101/2020.09.25.312793

**Authors:** Burcu Firatligil-Yildirir, Gizem Bati-Ayaz, Ismail Tahmaz, Muge Bilgen, Devrim Pesen-Okvur, Ozden Yalcin-Ozuysal

## Abstract

Metastasis is one of the major obstacles for breast cancer patients. Limitations of current models demand the development of novel platforms to predict metastatic potential and homing choices of cancer cells. Here, two novel Lab-on-a-chip (LOC) platforms, invasion/chemotaxis (IC-chip) and extravasation (EX-chip), are presented for the quantitative assessment of invasion and extravasation, towards specific tissues. On IC-chip, invasive MDA-MB-231, but not non-invasive MCF-7 breast cancer cells invaded lung and liver microenvironments. Lung-specific but not bone-specific MDA-MB-231 clones efficiently invaded lung microenvironment, stressing ability of IC-chip to demonstrate different *in vivo* metastatic behaviors. On EX-chip, MDA-MB-231 cells extravasated more into the lung microenvironment compared to the liver and breast highlighting the potency of the platform to mimic *in vivo* homing choices. Overall, this study presents IC-chip and EX-chip that can determine tissue-specific invasion and extravasation potentials of cancer cells providing the groundwork for novel diagnostic tools to predict metastasis risk.

## Introduction

Metastasis is the main cause of breast cancer mortality among women. The latest statistics revealed that the 5-year relative survival rate for women with metastatic breast cancer is around 26% between 1975-2017 (Howlader et al., 2020). Breast cancer frequently metastasizes to bone, lung, brain and liver. Although the molecular and histopathological subtype of the tumor provide information on the metastasis risk and the target organs, there is no diagnostic tool available that can accurately predict the risk and the organ preference for an individual patient’s tumor. The target organ for metastasis is specified by both physiological architectures of the circulatory system and molecular determinants. It was first hypothesized by Stephan Paget in the 19^th^ century that metastasizing tumor cells grow preferentially in specific target organs in a similar manner that a “seed” grows only in a suitable “soil” (Paget, 1889). Since then, experimental evidence have been supporting the “seed and soil” hypothesis by showing that molecular determinants on primary tumor and microenvironment of the target tissue are involved in the establishment of metastasis patterns (Langley and Fidler, 2011). Thus, a platform that integrates the information coming from tumor cells and the target organs would provide a diagnostic tool that can estimate the likeness of metastasis for a given tumor cell population towards specific environments.

The metastasis cascade is a complex phenomenon that starts with invasion and migration, followed by intravasation that releases the tumor cells into the circulation. Then the circulating tumor cells extravasate to the target tissues for colonization. The invasion process starts when cancer cells dissociate from their primary sites after losing cell-cell adhesion capacity and invade the surrounding stroma, while extravasation process requires interaction between cancer cells and endothelial cells, which then leads cancer cells to penetrate the endothelial layer in the target organ (Fares et al., 2020). Here, we focus on invasion and extravasation steps of the cascade to estimate the metastatic potential of the cells towards different environments.

*In vivo* animal models provide a great physiological setup for the investigation of the metastatic process, yet they have several obstacles to track the dynamic process and cannot offer a diagnostic platform. The Boyden chamber and transwell systems are the most favored *in vitro* platforms to study invasion and extravasation due to their simplicity. However, they use membranes which are artificial barriers and they cannot provide detailed visualization of cellular behavior at multiple time points. On the other hand, Lab-on-a-chip (LOC) systems that refer to a miniature laboratory device that scales down to millimeter size present great advantages for 3D i*n vitro* strategies. LOC shows low-cost features, increases reproducibility and accelerates the examination’s strength and stability. LOC platforms can mimic physiological conditions both at tissue and organ levels and therefore they have a huge potential to minimize of the requirement for animal testing in the preclinical research area (Soscia et al., 2017).

*In vitro* LOC models were developed to investigate different factors involved in metastasis such as intravasation, (Song et al., 2009, Truong et al., 2016) angiogenesis, (Vickerman and Kamm, 2012) the interaction between tumor cells and endothelial cells with stromal cells, (Zervantonakis et al., 2012) the interstitial flow, (Polacheck et al., 2011) matrix stiffness, (Pathak and Kumar, 2012) and extravasation.

Here we developed two LOC platforms to quantitatively investigate the invasion/chemotaxis and extravasation of cancer cells towards different microenvironments relevant to the organ-specific metastases (Bersini et al., 2014, Chen et al., 2017, Jeon et al., 2015).

## Results

### Design of and procedure for invasion-chemotaxis and extravasation chips

Invasion and extravasation are two of the crucial steps in cancer metastasis. To investigate tissue-specific invasion and extravasation, two different lab-on-a-chip (LOC) devices were used (Figure 1). The IC (invasion/chemotaxis) chip comprised three channels: media channel 1 (MC1), homing matrix channel (HMC), and media channel 2 (MC2). The IC-chip was symmetric along the long axis of HMC so that a gradient of factors in the MC2 can be realized across the HMC from MC2 to MC1. The procedure for invasion assay on the IC-chip is shown in Figure 1a. Here, cells loaded into the MC1 were expected to show invasion and chemotaxis in response to the microenvironment in the HMC and/or the contents of the MC2.

**Figure 1.**
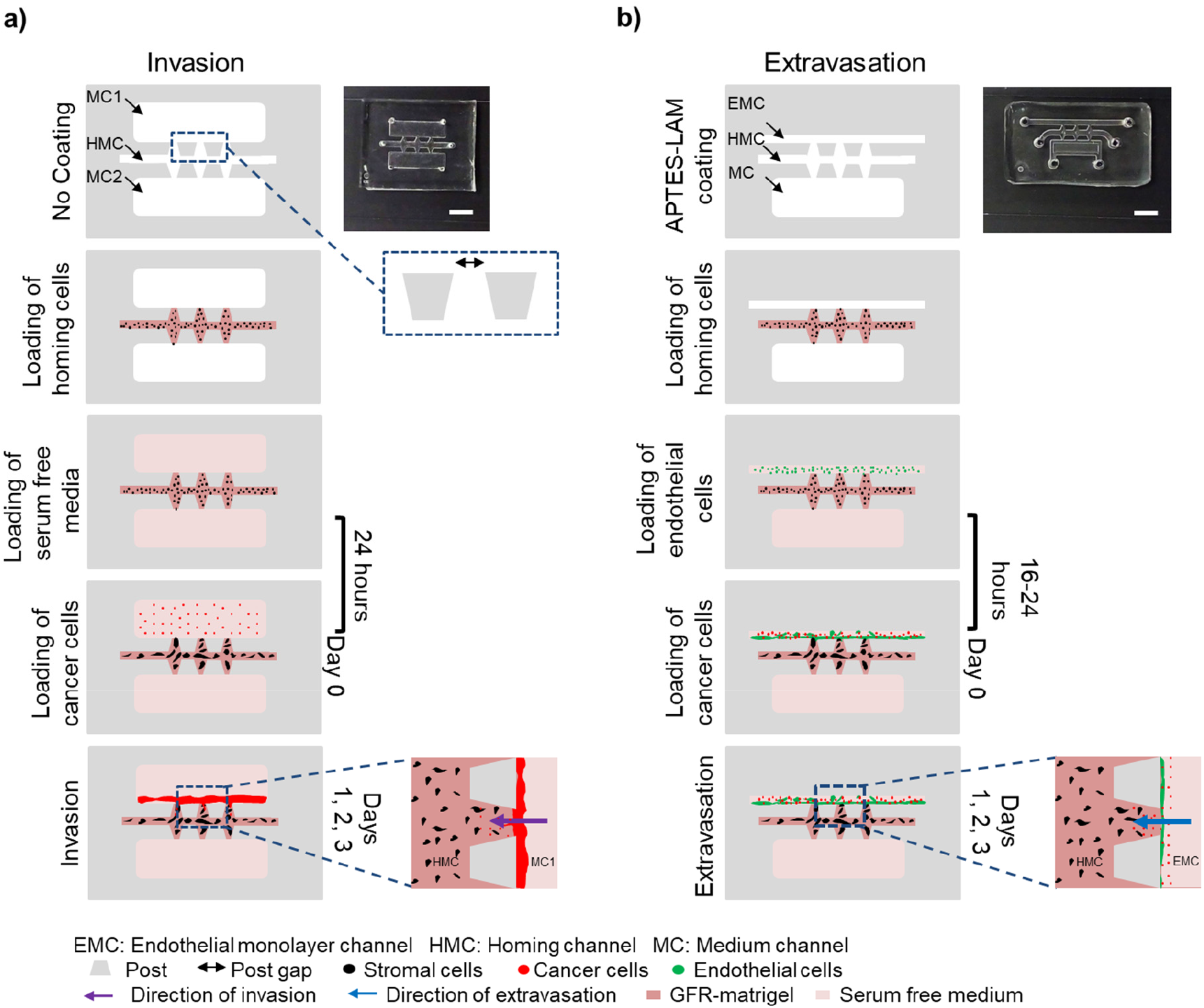
LOCs’ setup for invasion and extravasation assays. a) Schematic representation of invasion assay steps (loading of homing cells, loading of serum-free media, loading of cancer cells after 24 hours, and tracking of invasion on day 1, day 2 and day 3. On the left, a photo of invasion LOC and zoom-in images of post gaps and invasion from MC1 into HMC are shown). b) Schematic representation of extravasation assay steps (APTES-Lam coating of EX-chips, loading of homing cells and loading of endothelial cells in serum-free media, loading of cancer cells after 16-24 hours, tracking of extravasation for day 1, day 2 and day 3. On the left, a photo of extravasation LOC and zoom-in image of extravasation from EMC into HMC are shown). (EMC: Endothelial monolayer channel, HMC: Homing Channel, MC: Medium Channel, Stromal cells: Black, Endothelial cells: Green, Cancer cells: Red, Serum-free media: Light Pink, Growth Factor Reduced (GFR)-matrigel: Dark Pink, PDMS and Posts: Grey, Post gap: Black two-sided arrow, The direction of invasion: Purple arrow, The direction of extravasation: Blue arrow) (Scale bar: 5 mm)

The EX (extravasation) chip also comprised three channels: endothelial monolayer channel (EMC), homing matrix channel (HMC), medium channel (MC). However, the EMC and the MC were not mirror images of each other as was the case for the MC1 and MC2. The EMC was a narrow channel designed to hold endothelial cells and intended to mimic a blood vessel. The procedure for extravasation assay using the EX-chip is shown in Figure 1b. Here, cells loaded into the EMC after the formation of an intact endothelial monolayer were expected to show extravasation through the endothelial cells in response to the microenvironment in the HMC.

In both the IC-chip and the EX-chip, the HMC was used to mimic the microenvironments of different tissues such as lung, liver and breast. Thus, choices of invasion-chemotaxis and extravasation into different tissues can be investigated.

### Effect of serum on the invasion of breast cancer cells in the presence and absence of homing cells

Invasion and chemotaxis of tumor cells towards various factors in their microenvironment is one of the crucial steps of cancer metastasis. The factors secreted from stromal cells residing within target tissue microenvironments play important roles in chemotactic behavior of tumor cells towards their specific target sites (Fares et al., 2020, Guo and Deng, 2018, Roussos et al., 2011). Invasion-chemotaxis chip (IC-chip) was optimized as a tool for the analysis of cellular invasion and chemotaxis. Cell-free matrigel was loaded into the homing channel (HMC). Medium channel 2 (MC2) was loaded with culture media with or without 10% fetal bovine serum (FBS). Medium channel 1 (MC1) was loaded with triple-negative, invasive MDA-MB-231 cells stably labelled with DsRed and resuspended in serum-free media. Invasion and chemotaxis were observed for 3 days with confocal microscopy (Figure 2a).

**Figure 2.**
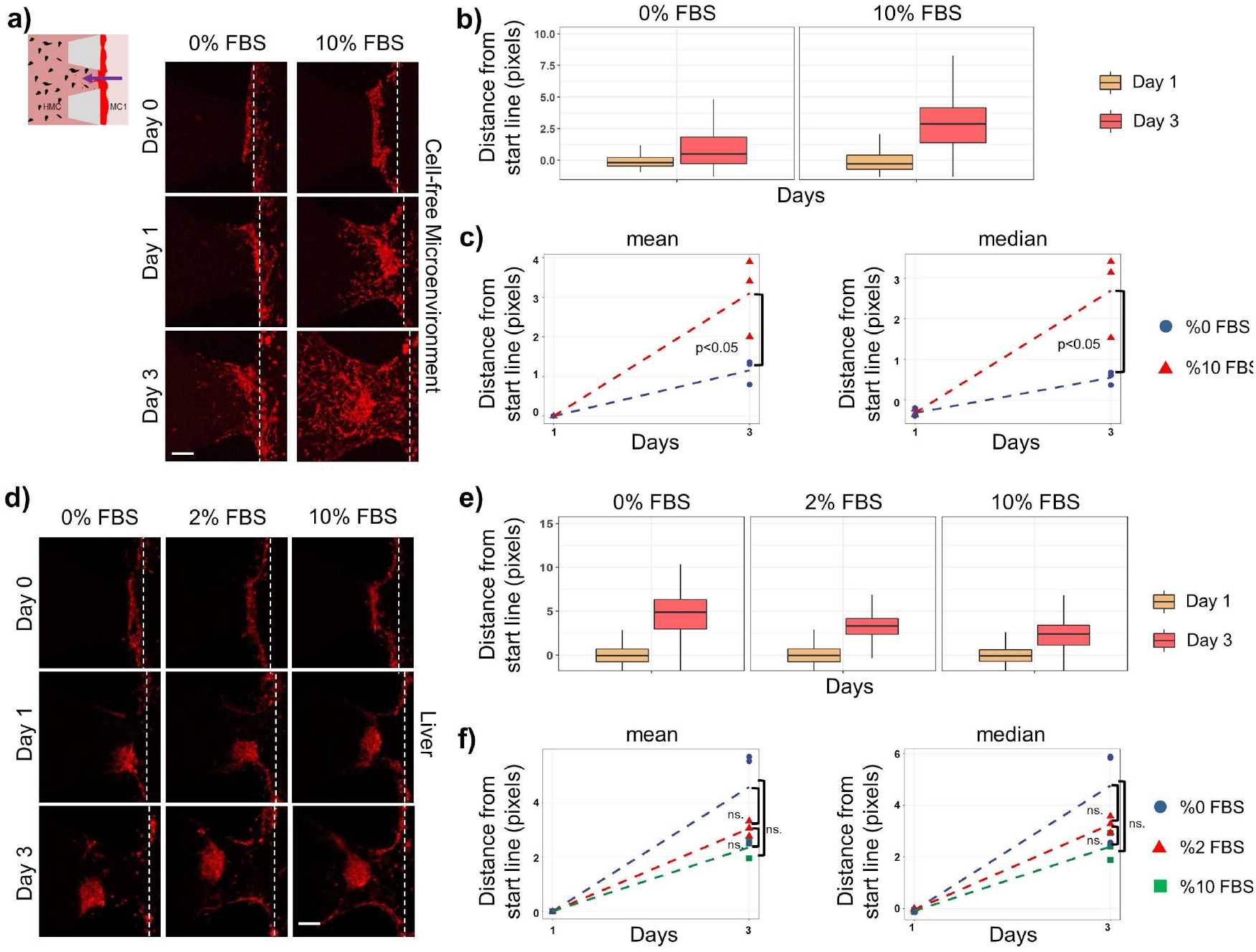
Invasion of MDA MB 231 cells towards empty matrigel and liver microenvironment in the presence or absence of serum. Representative Z-stack images showing invasion of MDA-MB-231 cells (red) towards a) empty matrigel in the absence (0%) or presence (10%) of serum and schematic representation of invasion from MC1 into HMC; and d) liver environment generated by BRL-3A with different serum concentrations (0, 2 and 10%) (dashed line corresponds to the starting line for invasion) (Scale bar: 200 μm). b) and e) The distance of each bright pixel to the starting line (dashed) was calculated after thresholding of Z-stack images. The data normalized to day 1 were plotted (n=3). c) and f) Mean and median values of normalized distance distributions were plotted for day 1 and day 3 (n=3). (FBS: fetal bovine serum)

Quantitative analysis of fluorescence images showed that invasion-migration increased from day 1 to day 3 for both FBS (10%) and FBS-free conditions, consistent with the invasive phenotype of MDA-MB-231 cells. (Figure 2b). However, MDA-MB-231 cells showed more invasion-migration towards FBS (10%) containing media than towards FBS free medium (p<0.05), as expected. A significant increase was detected in both the mean and the median distances invaded by the cells in FBS (10%) condition compared to the FBS-free condition (Figure 2c). Thus, IC-chip was able to demonstrate the chemoattractant role of serum visually and quantitatively.

To optimize the invasion conditions towards different tissue microenvironments, the effect of serum in the presence of homing cells was tested. BRL-3A liver cells were resuspended in serum-free media, mixed with GFR-matrigel and loaded into the HMC. Medium (0%, 2% or 10% FBS) was added into the MC2. Invasion of MDA-MB-231 cells loaded into the MC1 towards the HMC was examined (Figure 2d). Quantitative image analysis showed that for all the three different FBS concentrations in the MC2, the invasion of MDA-MB-231 cells increased from day 1 to day 3 (Figure 2e). Besides, there were no significant differences in the invasion of MDA-MB-231 cells with (2% or 10%) or without (0%) FBS containing media in the MC2 in the presence of liver cells in the HMC. The mean and the median distances invaded by the cells were not significantly different in serum-free condition (0%) compared to the serum containing media (2% and 10%) (Figure 2f). Therefore, the presence of homing cells such as BRL-3A was sufficient to induce invasion and chemotaxis of MDA-MB-231 cells. Consequently, serum-free media was used in the MC2 for all invasion experiments with cell-laden matrigel in the HMC. Altogether these results demonstrated that the IC-chip provided a robust platform for invasion assays either with or without target tissue cells.

### Invasion of breast cancer cells into the lung, liver and breast microenvironments

Invasion capacity of MDA-MB-231 and MCF-7 cells into lung, liver and breast microenvironments generated by WI-38, BRL-3A and MCF-10A cells embedded in GFR-matrigel in the HMC, respectively, was investigated (Figure 3a).

**Figure 3.**
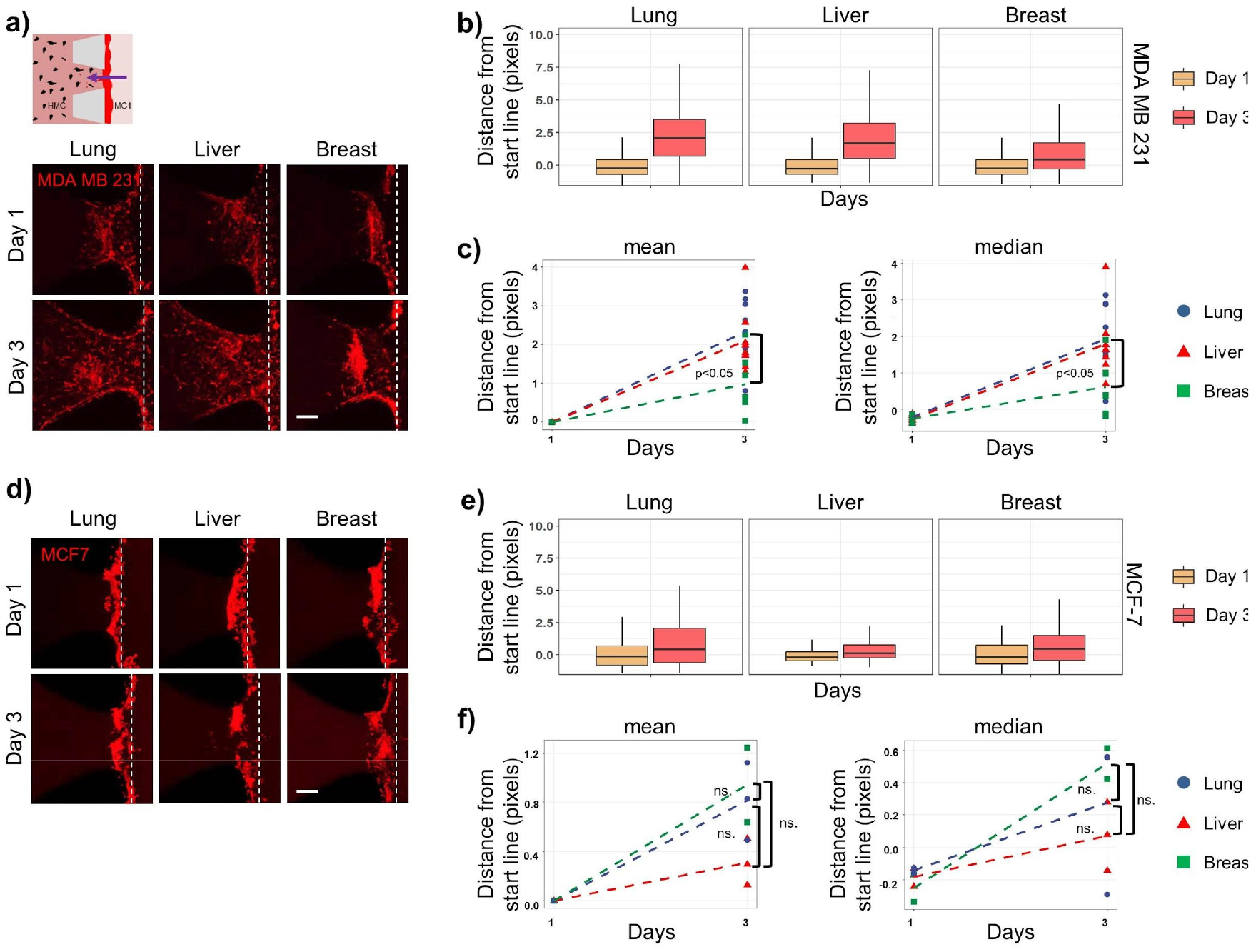
Invasion of metastatic and non-metastatic breast cancer cells towards lung, liver and breast microenvironments. Representative Z-stack images showing invasion of a) MDA-MB-231 (red) and d) MCF7 cells (red) towards lung, liver and breast microenvironments generated by WI-38, BRL-3A and MCF10A cells, respectively and a) schematic representation of invasion from MC1 into HMC (dashed line corresponds to the starting line for invasion) (Scale bar: 200 μm). b) and e) The distance of each bright pixel to the starting line (dashed) was calculated after thresholding of Z-stack images. The data normalized to day 1 were plotted (n=6). c) and f) Mean and median values of normalized distance distributions were plotted for day 1 and day 3 (n=6).

The distance MDA-MB-231 cells invaded towards all the three microenvironments increased from day 1 to day 3 (Figure 3b). However, the invasion of MDA-MB-231 cells to lung and liver microenvironments was significantly higher than that to the breast microenvironment (Figure 3c and Table S1, Supporting Information). Non-metastatic MCF-7 cells did not significantly invade towards lung, liver or breast environments up to day 3. (Figure 3d, e, f). These results showed that MDA-MB-231 cells had a higher preference of invasion to the lung and liver microenvironments than the breast microenvironment, as expected, while the non-invasive MCF7 cells did not invade and thus had no preference for different homing tissues.

### Invasion of lung-specific and bone-specific metastatic breast cancer cells into the lung microenvironment

Lungs are the most common site of breast cancer metastasis (Jin et al., 2018). Here, we examined the invasive potential of organ-specific metastatic clones of MDA-MB-231 cells for lung (MDA-MB-231 LM2) and bone (MDA-MB-231 1833-BoM) (Bos et al., 2009, Kang et al., 2005, Minn et al., 2005) towards lung microenvironment in the IC-chip. Parental and lung-specific (LM2) MDA-MB-231 cells invaded the lung microenvironment remarkably well, while bone-specific (BoM 1833) cells moved marginally towards HMC (Figure 4a, b). The distance invaded by parental and lung-specific parental cells were significantly longer than bone-specific parental cells (Figure 4c and Table S1, Supporting Information). These data suggest that the lung microenvironment generated by WI-38 cells embedded in GFR-matrigel in the IC-chip successfully mimicked the *in vivo* lung microenvironment.

**Figure 4.**
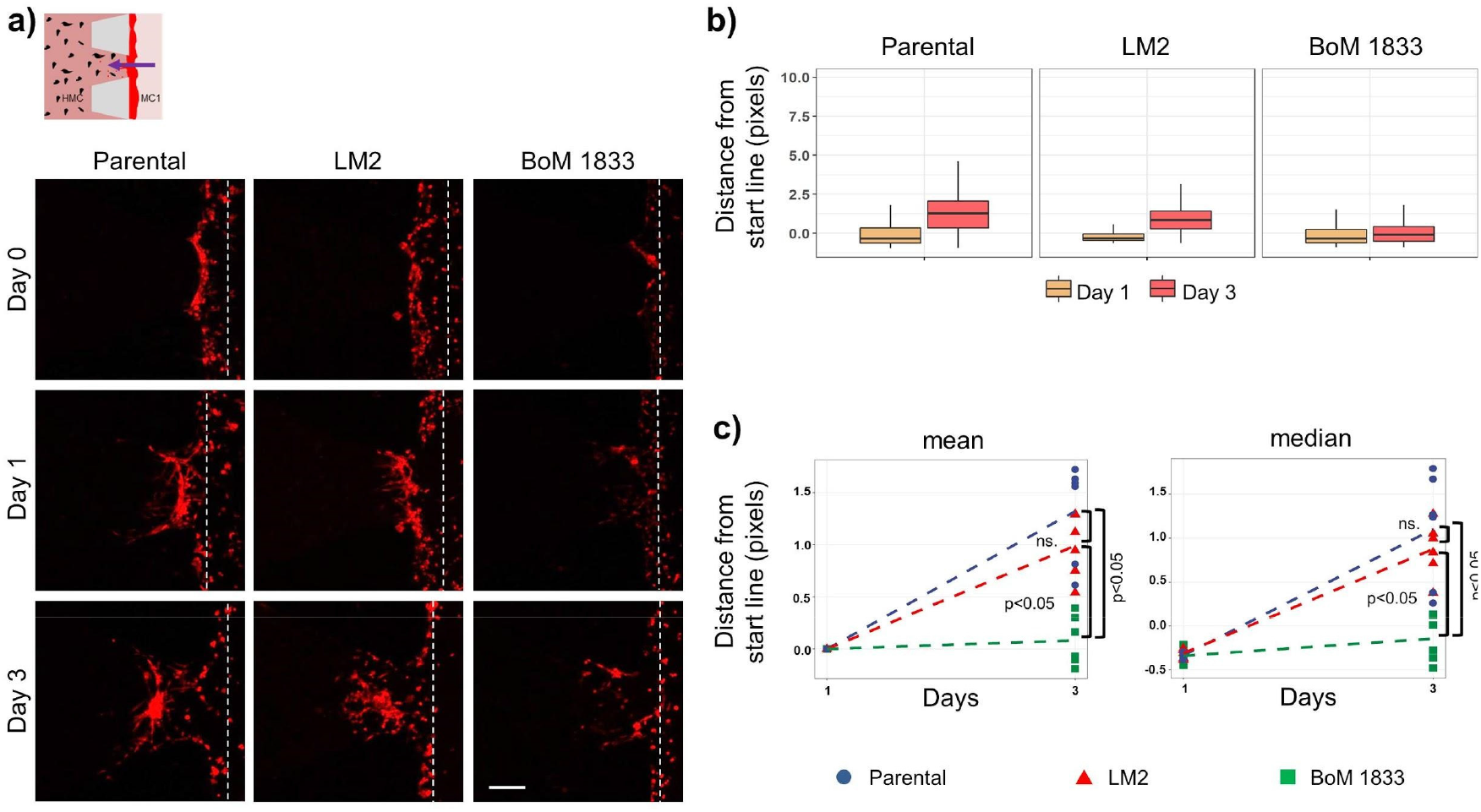
Invasion of lung-specific and bone-specific MDA-MB-231 clones towards the lung microenvironment. a) Representative Z-stack images showing invasion of parental, LM2 (lung-specific) and BoM 1833 (bone-specific) MDA-MB-231 cells (red) towards the lung microenvironment generated by WI-38 cell line (dashed line corresponds to the starting line for invasion) and schematic image of invasion from MC1 into HMC (Scale bar: 200 μm). b) The distance of each bright pixel to the starting line (dashed) was calculated after thresholding of Z-stack images. The data normalized to day 1 were plotted (n=3). c) Mean and median values of normalized distance distributions were plotted for day 1 and day 3 (n=3).

### Generation of the intact endothelial monolayer

An intact endothelial monolayer is essential for mimicking extravasation. When endothelial cells were loaded into EMC, they tended to form clusters. Here, dextran was used in the cell resuspension medium to inhibit cluster formation and ensure a homogeneous distribution of endothelial cells in the EMC (Figure S1a, Supporting Information) (Myers et al., 2012).

Before the extravasation assays, the interior surfaces of EX-chips were chemically and biochemically modified to ensure the attachment of endothelial cells for the generation of an intact monolayer. Here, *3-Aminopropyl triethoxysilane* (APTES) and Poly-L-lysine solution (PLL) were tested for their ability to promote fibronectin (FN) coating. Both APTES and PLL supported FN coating and thus efficient endothelial cell monolayer formation. (Figure S1b, Supporting Information).

APTES coating was preferred due to the shorter application time. Formation of an intact endothelial monolayer is a critical step for mimicking blood vessels in extravasation assays. To optimize the formation of an intact endothelial monolayer, laminin (LAM), collagen type I (COL) and fibronectin (FN) were tested on APTES pre-coated interior PDMS surfaces of EX-chips. LAM coated surfaces provided the most appropriate surfaces for the attachment of endothelial cells that covered a larger area (Figure 5a, b).

**Figure 5.**
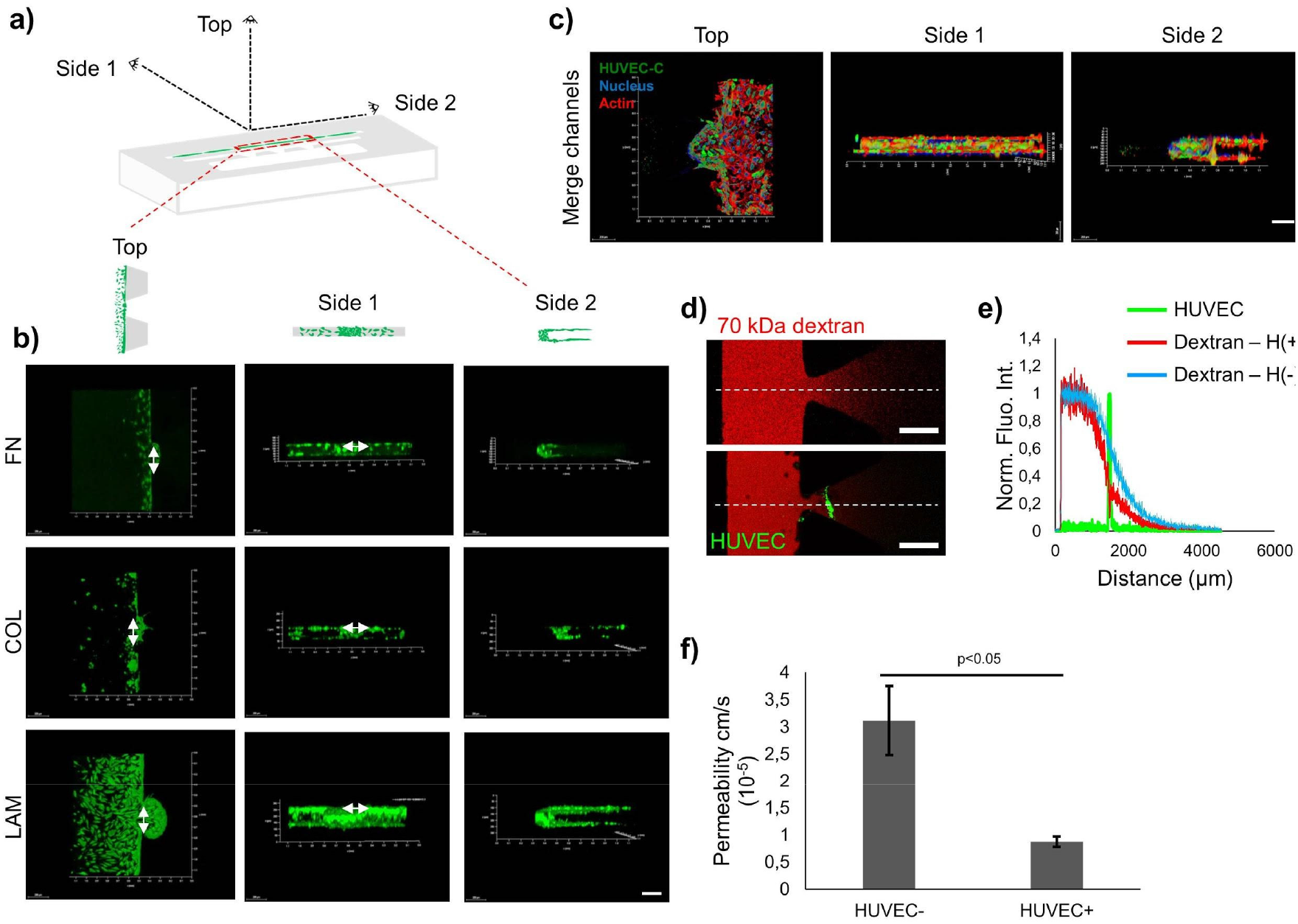
Laminin coating resulted in more efficient intact endothelial monolayer formation. a) Schematic of 3D Ex-chip from different views (Top, Side1, Side 2). b) Representative 3D images showing endothelial cells (green) on fibronectin (FN), collagen type I (COL), and laminin (LAM) coated surfaces between the post gaps are represented with two-sided arrows from different views (Scale bar: 200 μm). c) Confocal images showing actin (phalloidin), nuclei (DAPI), HUVEC-C (cell tracker) with merge channels in red, blue and green, respectively from different views in APTES-LAM coated EX-chip (Scale bar: 200 μm). d) A representative image of 70 kDa fluorescent dextran (red) in a 3D homing channel (HMC) region in the absence (top panel) and the presence (bottom panel) of the endothelial monolayer (green) (Scale bar: 500 μm). e) Normalized fluorescent intensity profiles along the dashed line. HUVEC-C signal (green), dextran signal in the absence (blue) and presence (red) of HUVEC-C. f) Diffusive permeability of cell-free GFR-matrigel and endothelial monolayer for 70 kDa dextran.

Intact endothelial monolayer formation was confirmed by staining the cytoskeleton of endothelial cells, specifically, actin filaments (Figure 5c). Fluorescence signal coming from Green Cell Tracker labelled HUVEC-C cells was sparse. However, actin staining confirmed the confluence of the endothelial monolayer. These results suggested that the green signal obtained by transient labelling of cells by the Green Cell Tracker might not reflect the extent of HUVEC-C coverage on the surface. Intact endothelial monolayer formation was further demonstrated by measuring diffusion of fluorescent 70-kDa dextran from the EMC to the HMC in the presence and absence of endothelial cells (Figure 5d, e) Permeability calculations showed that presence of an endothelial monolayer significantly reduced diffusion of fluorescent 70 kDa dextran from 3.12 +/-0.63 × 10^−5^ cm s^-1^ to 0.88 +/-0.1 × 10^−5^ cm s^-1^ (p-value < 0.02) (Figure 5f). Altogether, these results demonstrated that an intact endothelial monolayer with low permeability can be achieved by using APTES-laminin coating in EX-chips.

### Extravasation of metastatic breast cancer cells into the lung, liver and breast microenvironments

To analyze the homing choices of extravasating breast cancer cells, lung, liver and breast microenvironments were generated in the HMC, while an intact endothelial monolayer was generated in the EMC of the EX-chip (Figure 6a, b). Extravasation metric (EM) is a measure of the efficiency and reproducibility of the EX-chips. EM for the lung, liver and breast microenvironments were 0.89, 1 and 0.89, respectively, with no statistically significant differences between different microenvironments, showing that extravasation events were observed in almost all post gaps independent from the homing microenvironment. Cancer cells that passed through the endothelial layer were considered as extravasated, while cells that were detected within the endothelial layer were considered as associated. Extravasated cell numbers were from highest to lowest in the lung, liver and breast microenvironments (Figure 6c and Table S1, Supporting Information). Numbers of MDA-MB-231 cells that remained associated with the endothelial monolayer were highest where the HMC contained the breast microenvironment (Figure 6c and Table S1, Supporting Information).

**Figure 6.**
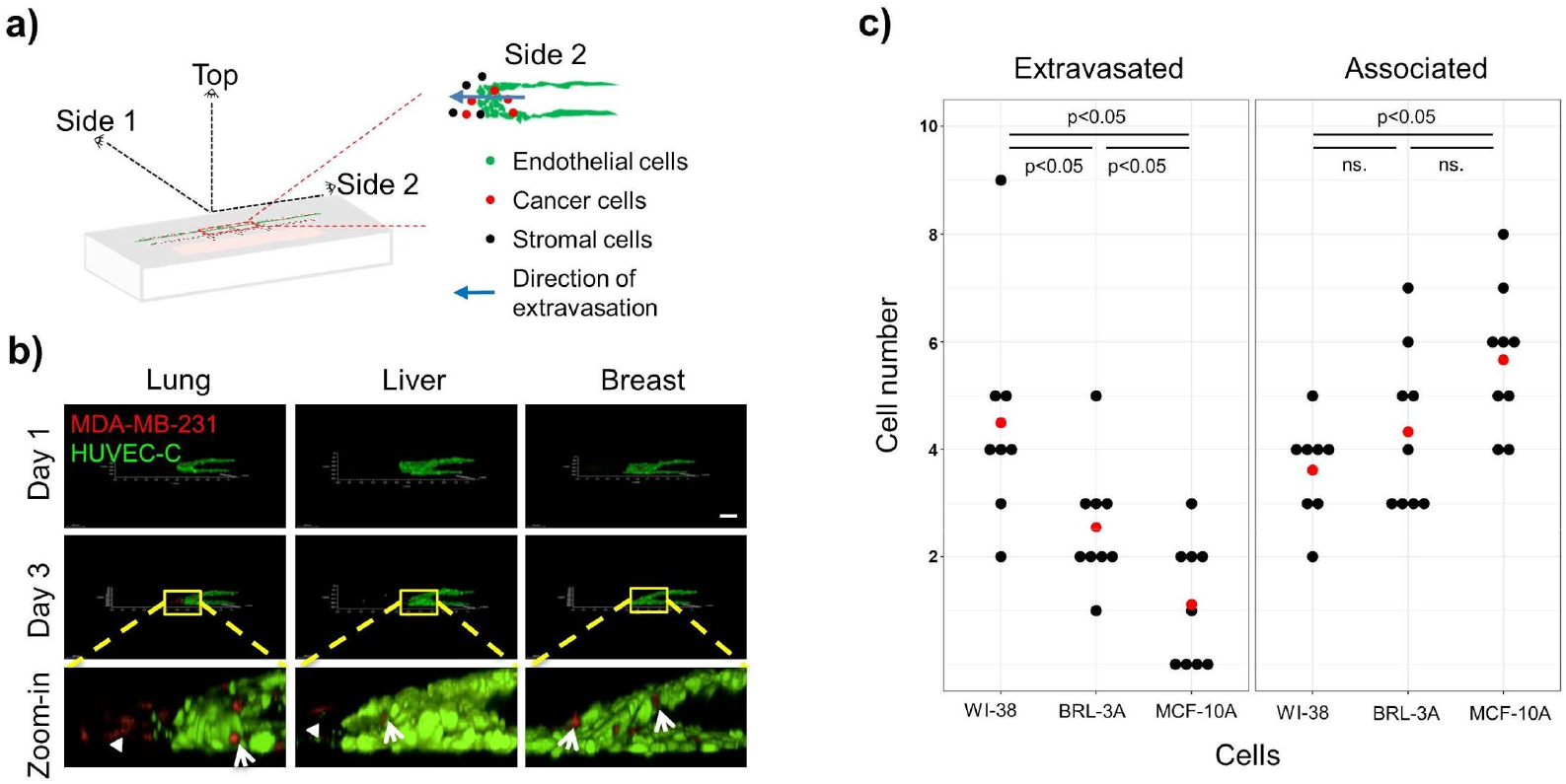
Extravasation of metastatic breast cancer cells into lung, liver and breast microenvironments. a) 3D schematic of the EX-chip. Zoom-in image of 3D EX-chip showing the side 2 view (Cancer cells: Red, Endothelial cells: Green, Direction of extravasation: Blue arrow). b) Representative Z-stack images showing side 2 views of endothelial monolayer formation by HUVEC-C cells (green) and extravasated (arrow head) or associated (arrow) MDA-MB-231 cells (red) into lung, liver or breast microenvironments generated by WI-38, BRL-3A and MCF-10A cell lines, respectively (Scale bar: 200 μm). c) The number of extravasated and associated MDA-MB-231 cells. Each black dot represents the cell number for one post gap within the EX-chip, while the red dot is the average number of cells for each condition (n=9).

## Discussion

Microfluidic platforms such as lab-on-a-chip (LOC) systems are being studied in several research areas from electronics to biology and they currently take part in a crucial number of studies focusing on metastasis process including metastatic breast cancer. Therefore, it is prominent to develop a LOC system providing a convenient platform for real-time monitoring of the cancer cell behavior involved in the metastatic cascade. In the present study, we introduced two novel LOC systems that enable the visualization and quantification of tissue-specific breast cancer invasion and extravasation. In this context, we generated lung, liver and breast microenvironments as potential homing tissues for metastatic breast cancer cell line, MDA-MB-231, and tested the invasion and extravasation potential of cancer cells towards these target sites.

The generation of endothelial monolayer is a critical step for metastasis research to mimic blood vessels. This approach mainly consists of the attachment and distribution of endothelial cells. Due to the hydrophobic nature of cured PDMS that does not allow a convenient surface for cell attachment, surface modifications are needed such as APTES and PLL coatings (Leivo et al., 2017). In our approach, APTES was used as a linker chemical to promote protein attachment. Endothelial cells lining blood vessels act as a barrier to cell movement that should be impaired or remodeled by cancer cells for their extravasation (Shenoy and Lu, 2016). To optimize the formation of an intact endothelial monolayer, laminin, which is a component of the basement membrane, collagen type I, which is abundant in connective tissue and FN, a common extracellular matrix protein, were tested on the APTES pre-coated interior PDMS surfaces of EX-chips. Among the three different proteins, laminin promoted endothelial cell adhesion most. This is probably because endothelial cells *in vivo* exist on a basement membrane, of which laminin is a marked component. The endothelial barrier function was quantified to confirm the size-selective transendothelial transport of cancer cells. The 70 kDa dextran used here to measure endothelial monolayer integrity is equal to the molecular weight of serum albumin, which is 69 kDa (Ono et al., 2005). The permeability of the endothelial monolayer generated in our microfluidic platform with and without endothelial cells are in agreement with the previous studies (Frost et al., 2019, Jeon et al., 2013, van Duinen et al., 2017, Zervantonakis et al., 2012).

The metastasis of triple-negative breast cancer is mainly seen in the lung and liver (Al-Mahmood et al., 2018). Consistent with the clinical data, results here showed that MDA-MB-231 cells, which fall into the category of triple-negative subtype, preferred lung and liver microenvironments over the breast microenvironment. MDA-MB-231 cells invading towards the lung and liver microenvironments showed dispersed organization, single-cell migration and longer invasion distances whereas MDA-MB-231 cells invading towards the breast microenvironment appeared to move in a sheet-like manner, suggesting collective migration (Friedl and Wolf, 2003). The preferential dissemination of MDA-MB-231 cells to the lungs has been shown (Chu et al., 2014, Minn et al., 2005). Here the representative lung microenvironment generated by WI-38 cells to mimic *in vivo* conditions was tested with lung metastatic subclone MDA-MB-231 LM2, which preferred the lung microenvironment, consistent with previous *in vivo* studies. Parental MDA-MB-231 cells also showed a similar trend in the invasion towards lung environment as seen in MDA-MB-231 LM2, but not in bone-specific MDA-MB-231 1833-BoM suggesting that the lung-specific metastatic cells could be more populated in the parental MDA-MB-231 cell line. Taken together, these results highlight the ability of the IC-chip to quantitatively determine the invasion potential of cancer cells towards different target sites and to distinguish between cancer cells with different *in vivo* metastatic behaviors.

Finally, both the invasion and extravasation assays showed that MDA-MB-231 cells preferred the lung microenvironment significantly more than the other target microenvironments. Although the MDA-MB-231 cells invaded towards the breast microenvironment on IC-chip, they did not extravasate into the same microenvironment. These findings indicate that IC-chip would be useful to predict invasive behavior of the cancer cells in the primary tumor site as well as their preference for different target tissues. On the other hand, EX-chip would be more relevant and efficient for the determination of the overall metastatic potential and the homing choices of breast cancer cells.

In conclusion, we developed two LOC platforms, IC-chip and EX-chip, to quantitatively investigate the invasion/chemotaxis and extravasation of cancer cells towards different microenvironments. We tested the LOC platforms with invasive MDA-MB-231 and non-invasive MCF-7 breast cancer cells. The observations we made were consistent with the *in vivo* behavior of the cells that MDA-MB-231 cells specifically invaded and extravasated towards lung and liver microenvironments, while MCF-7 cells did not show an invasive phenotype. The IC-chip and the EX-chip platforms developed here lay the basis for the development of diagnostic LOC devices to quantify the invasive behavior and tissue-specific extravasation potential of cancer cells to predict metastasis risk for breast cancer patients.

## Materials and Methods

### Lab-on-a-chip fabrication

Invasion-chemotaxis and extravasation lab-on-a-chip platforms (IC-chip and EX-chip) were either provided by Initio Biomedical Engineering (Turkey) or fabricated by soft lithography as previously described (Ozdil et al., 2014). Briefly, SU-8 polymer was spin-coated on a silicon wafer. The design of the chip was exposed through a mask to UV light. After removing the uncrosslinked SU8 polymer using the developer solution, molds were ready for polydimethylsiloxane (PDMS) casting. After PDMS polymerization, inlet and outlet holes were punched with biopsy punches. The PDMS parts were cleaned and bonded to clean microscope slides after treatment in UV/Ozone cleaner (Bioforce Nanosciences, USA). The chips were sterilized with UV light in a laminar hood for 15 minutes before use. The dimensions of IC-chip: the homing matrix channel (HMC) 3 mm width x 12 mm length x 200 μm height and medium channels (MC1/MC2) 3 mm width x 12 mm length x 200 μm height, EX-chip: the homing matrix channel (HMC) 3 mm width x 15 mm length x 200 μm height, endothelial monolayer channel (EMC) 3 mm width x 20 mm length x 200 μm height and medium channel (MC) 3 mm width x 10 mm length x 200 μm height.

### Chip surface modification

IC-chips were used without any surface modifications. EX-chips were first coated with either poly-l-lysine (PLL, P8920, Sigma Aldrich) or *3-aminopropyltriethoxysilan*e (APTES, A3648, Sigma Aldrich). EX-chips were incubated with PLL (0.01 mg mL^-1^) in ultra-pure water at 37°C in a 5% CO_2_ incubator overnight. The following day, EX-chips were washed with ultra-pure water 3 times and then kept at 80°C for 24 hours to reduce hydrophobicity of the surface. For APTES modification of surfaces, APTES (2%) dissolved in acetone was loaded into the channels of EX-chips and incubated for 15 minutes in laminar flow cabin. Then EX-chips were washed with first PBS once and then ultra-pure autoclaved H2O thrice. At this step, EX-chips were ready to be coated with extracellular matrix proteins: laminin (LAM), type I collagen (COL) or fibronectin (FN). LAM (0.0125 mg mL^-1^, L2020, Sigma Aldrich) and FN (0.0125 mg mL^-1^, F2006, Sigma Aldrich) were prepared in 1X Universal Buffer (UB), while COL (0.0125 mg mL^-1^, 354249, Corning) was diluted in serum-free DMEM (Biological Industries, 01-055-1A). Each protein solution was loaded into EX-chips and they were incubated at 37°C in a 5% CO_2_ incubator for 1 hour. LAM and FN coated EX-chips were washed first with PBS once and then with ultra-pure autoclaved H2O thrice. COL coated EX-chips were washed with first serum-free DMEM and then ultra-pure autoclaved H2O for thrice. Any remaining H2O was aspirated by vacuum. EX-chips were stored in vacuum desiccators at least one day before use in experiments. APTES-LAM coated EX-chips were used in all extravasation assays (Figure 1b).

### Cell Lines

Human breast cancer cell lines (MDA-MB-231 and MCF-7), human normal mammary epithelial cell line (MCF-10A), human normal lung fibroblast cell line (WI-38), rat normal liver cell line (BRL-3A), and human umbilical vein endothelial cell line (HUVEC-C) were obtained from ATCC. Organ-specific metastatic clones of MDA-MB-231, MDA-MB-231 LM2 and MDA-MB-231 1833-BoM were described previously (Bos et al., 2009, Kang et al., 2005, Minn et al., 2005) and were gifts from the Joan Massagué Lab in Memorial Sloan Kettering Cancer Center. MDA-MB-231, its derivatives and MCF-7 were cultured in DMEM high glucose (11965092, Gibco) with Fetal Bovine Serum (FBS, 10%) (A3840001, Gibco) and Penicillin/Streptomycin (15070063, Gibco, 1%); MCF-10A was cultured in DMEM-F12 high glucose (11330057, Gibco) with Horse Serum (04-004-1A, Biological Industries, 5%), Insulin (I9278, Sigma, 10 μg mL^-1^), Cholera Toxin (C8052, Sigma, 100 ng mL^-1^), EGF (E9644, Sigma, 20 ng mL^-1^), Hydrocortisone (H0888, Sigma, 0.5 μg mL^-1^) and Penicillin/Streptomycin (1%). WI-38 and BRL-3A were cultured in high glucose MEM-α (01-042-1A, Biological Industries) with Fetal Bovine Serum (FBS, 10%) and Penicillin/Streptomycin (1%). HUVEC-C cell line was cultured in DMEM-F12K high glucose (01-095-1A, Biological Industries) with Fetal Bovine Serum (FBS, 10%), Heparin (H3393, Sigma, 0.1 mg mL^-1^), endothelial cell growth supplement (EGCS, 0.05 mg mL^-1^) (354006, Sigma) and Penicillin/Streptomycin (1%). All cell lines were cultured at 37°C in a humidified incubator with 5% CO_2_.

### Labelling of Cell Lines

MDA-MB-231, metastatic clones of MDA-MB-231 (LM2 and 1833-BoM) and MCF-7 cancer cell lines were stably labelled with a red fluorescent protein (DsRed). MSCV retroviruses expressing both DsRed and puromycin resistance genes were used for infection. The preparation of viruses and infection of cells were performed as described previously. (Yalcin-Ozuysal et al., 2010, Zengin et al., 2015) 48 hours after infection, the antibiotic selection was carried out with puromycin (2 µg mL^-1^) until all of the uninfected cells died. Transient labelling of HUVEC-C cells was performed by Green Cell Tracker CMFDA (C2925, Invitrogen). The dye was dissolved in DMSO to obtain stock solution (25 mM) which was then diluted with serum-free DMEM-F12K media to get working concentration (5 µC). Cells were washed with warm PBS once, and then tracker (5 µC) was added over the cells. After 30 minutes of incubation at 37°C, the media was removed, cells were washed with PBS once and then complete HUVEC-C growth media was added. Labelling was performed 30 minutes before the experimental set-up.

### Invasion Assay

IC-chips were used for invasion assays (Figure 1a). For cell-free assays, growth factor reduced matrigel (GFR-matrigel, 8 mg mL^-1^) (354230, Corning) was diluted in 1:1 ratio with pre-cooled serum-free media and loaded into the homing matrix channel (HMC) of the chips. Then, chips were incubated for polymerization at 37°C in a humidified incubator with 5% CO_2_ for 30 minutes. After polymerization of GFR-matrigel, either serum-free or serum-containing media was loaded into the media channels 1 and 2 (MC1, MC2) and chips were incubated overnight. The following day, media in MC1 and MC2 were removed, channels were washed with serum-free media twice. Then serum-free (0%) or serum-containing media (10%) was added to MC2 of the chip for the relevant conditions. DsRed labelled MDA-MB-231 cells (1×10^6^ cells mL^-1^) resuspended in serum-free media were added to MC1. The chips were incubated vertically for 3 days.

To analyze the effects of serum on invasion towards liver microenvironment, BRL-3A normal liver cells (1×10^7^ cells mL^-1^) with GFR-matrigel were loaded to HMC of the IC-chips as explained above. Then, chips were incubated overnight with culture media with (2% or 10%) or without (0%) serum at both MC1 and MC2 channels. The following day, MC2 was loaded with serum-free media after washing the channel with serum-free media twice. DsRed labelled MDA-MB-231 cells (1×10^6^ cells mL^-1^) in serum-free media, were added to the MC1 and incubated vertically for 3 days.

To analyze invasion towards specific tissues, lung, liver and breast microenvironments were modelled by WI-38, BRL-3A and MCF-10A cells, respectively. Each cell line (BRL-3A: 1×10^7^ cells mL^-1^, WI-38: 5×10^6^ cells mL^-1^, MCF-10A: 4,4×10^6^ cells mL^-1^) was mixed with GFR-matrigel and loaded into the HMC of the IC-chips. The chips were incubated overnight with serum-free media in MC1 and MC2. The following day, media in MC2 was changed with fresh serum-free media. MDA-MB-231 or MCF-7 cells (1×10^6^ cells mL^-1^) were seeded to MC1 in serum-free media. Chips were incubated vertically for 3 days. The invasion was visualized every 24 hours by 3D imaging using a Leica SP8 confocal microscope.

### Analysis of Invasion

Z-stack images of IC-chips were acquired each day with a 10X objective and a z-step size of 7.52 µm. The analysis of the acquired images was performed by Python programming and R Studio as previously explained. (Ilhan et al., 2020) Briefly, the sum projection of z-stacks was thresholded and the distance of each bright pixel to the starting line of the invasion was calculated. The invasion capacity of the cells was determined through normalization of datasets to day 1.

### Endothelial monolayer formation

HUVEC-C cells labelled with Green Cell Tracker CMFDA were collected from culture dishes following Trypsin EDTA Solution A (0.25%, 03-050-1B, Biological Industries) treatment for 5 minutes. After centrifugation, they were resuspended in 450-650 kDa dextran (8%, 31392, Sigma Aldrich) in HUVEC-C media. The HUVEC-C cells (3.85x 10^6^ mL^-1^) were loaded to EMC of APTES-LAM coated EX-chips. EX-chips were incubated vertically at 37°C in a humidified incubator with 5% CO_2_ overnight. Endothelial monolayer formation was confirmed by 3D imaging using a Leica SP8 confocal microscope with a 10X objective and a z-step size of 7.52 µm.

### Actin staining for endothelial monolayer

Actin staining was performed to confirm the physical integrity of the endothelial cell monolayer. Cell-free GFR-matrigel (1:1 GFR-matrigel in serum-free media) was loaded to the HMC of the EX-chips. Following polymerization of matrigel, HUVEC-C cells were seeded to the EMC, culture media was loaded into the MC and chips were incubated overnight at 37°C in a humidified incubator with 5% CO_2_. The following day, media within EMC and MC was removed and paraformaldehyde (4%) was added to fix the sample. Then, the chip was incubated overnight at +4°C. The following day, the EMC and MC were washed with PBS (1X) thrice. Permeabilization solution (5% BSA and 0.1% Triton-X-100 in PBS buffer) was loaded to the EMC and MC and incubated at room temperature (RT) for 15 minutes. Then, the EMC and MC were washed with PBS (1X) thrice. Phalloidin (1:40, Alexa Fluor(tm) 647) (A22287, Invitrogen) for actin-filament staining and DAPI (1:500) for nuclei staining diluted in PBS were loaded to the EMC and MC and the chip was incubated for one hour at RT in the dark. Finally, the EMC and MC were washed with PBS and then filled with anti-fading mounting media (90% Glycerol, 10% PBS 10X, 0.1M or 2% (w/v) n-propyl gallate). The chip was kept at +4°C. The next day, images were acquired by a Leica SP8 confocal microscope (Figure 5c).

### Endothelial monolayer permeability assay

Fluorescently labelled 70-kDa dextran TR (D1830, Texas Red, neutral, Thermofisher) (final concentration 0.1 mg mL^-1^) was used for the measurement of endothelial monolayer permeability. 70-kDa dextran TR in PBS was loaded into the EMC. The chip was imaged using a Leica SP8 confocal microscope with 10X objective. Images were processed with ImageJ/Fiji and numerical analysis was performed using Excel. Data were processed as described previously.(van Duinen et al., 2017)

### Extravasation Assay

EX-chips were used for extravasation assays (Figure 1b). The same protocols for environment generation in the invasion assay and endothelial monolayer formation were followed as explained above. Once the monolayer was formed by HUVEC-C cells, DsRed labelled MDA-MB-231 cells (1×10^6^ cells mL^-1^) were seeded to the EMC in serum-free media for each condition (lung, liver and breast microenvironments) and the chips were incubated vertically for 3 days. The integrity of endothelial monolayer was confirmed by confocal microscopy immediately after addition of MDA-MB-231 cells. The extravasation of MDA-MB-231 cells to the generated lung, liver and breast microenvironments was visualized by 3D imaging using a Leica SP8 confocal microscope at 10x magnification and a z-step size of 7.52 µm for 3 days.

### Analysis of Extravasation

Region of interests (ROIs) were selected from the post gaps that comprising the interphase of EMC and HMC in the EX-chips. Only the ROIs, where intact endothelial monolayer was formed in the gap between two posts were included in the analysis. Cancer cells were marked as “extravasated” if they passed through the endothelial monolayer, or “associated” if they kept in contact with the endothelial monolayer. The efficiency of the EX-chips was quantified by the extravasation metric (EM), defined as the ratio of the number of post gaps with one or more extravasated cells to the total number of post gaps. If extravasation is observed in all post gaps, the EM will be 1. The χ2 (Chi-squared) test was used for the statistical analysis of the EM.

### Statistical Analysis

Results are reported mean ± s.e.m. unless otherwise noted. Student’s t-test was used for statistical analysis unless otherwise noted. A p-value of < 0.05 was considered significant.

## Acknowledgements

This work was supported by the grant numbered 115E057 from Scientific and Technological Research Council of Turkey (TUBITAK). The authors thank Joan Massagué and Memorial Sloan Kettering Cancer Center for providing MDA-MB-231 LM2 and MDA-MB-231 1833-BoM cell lines.

## Author Contributions

B.F.Y and G.B.A: Methodology, Formal Analysis, Investigation, Writing-Original Draft, Visualization, I.T and M.B: Methodology, Investigation, Formal Analysis, Writing-Original Draft, D.P.O and O.Y.O: Methodology, Conceptualization, Software, Formal Analysis, Resources, Writing-Original Draft, Wrirting-Review&Editing, Supervision, Project Administration, Funding Acquisition.

