## Supplementary Material for "Homing Choices of Breast Cancer Cells Revealed by Tissue Specific Invasion and Extravasation Lab-on-a-chip Platforms"

**Table S1.** Mean, median and p-values of the analysis shown in figures 3, 4 and 6.

| | Cancer Cells | Homing tissue (cells) | Mean ± s.e.m. | Median ± s.e.m. |
|---|---|---|---|---|
| Figure 3 | MDA-MB-231 | Lung (WI-38) | 2.33±0.27 | 1.94±0.31 |
| | | Liver (BRL-3A) | 2.10±0.30 | 1.80±0.33 |
| | | | **Mean p-value** | **Median p-value** |
| | | Lung vs Breast | 0.003 | 0.008 |
| | | Liver vs Breast | 0.014 | 0.018 |

| | Cancer Cells | Homing tissue (cells) | Mean ± s.e.m. | Median ± s.e.m. |
|---|---|---|---|---|
| Figure 4 | Parental MDA-MB-231 | Lung (WI-38) | 1.32±0.20 | 1.10±0.26 |
| | Lung specific (LM2) MDA-MB-231 | | 0.99±0.12 | 0.87±0.13 |
| | Bone specific (BoM 1833) MDA-MB-231 | | 0.008±0.10 | -0.14±0.11 |
| | | | **Mean p-value** | **Median p-value** |
| | | LM2 vs BoM 1833 | 4.85E-04 | 1.15E-03 |
| | | Parental vs BoM 1833 | 2.22E-05 | 1.95E-04 |

| | Cancer Cells | Homing tissue (cells) | # of extravasated cells | # of associated cells |
|---|---|---|---|---|
| Figure 6 | MDA-MB-231 | Lung (WI-38) | 4.50±0.73 | 3.63±0.33 |
| | | Liver (BRL-3A) | 2.56±0.38 | 4.34±0.50 |
| | | Breast (MCF-10A) | 1.11±0.39 | 5.67±0.44 |
| | | | **Extravasation p-value** | **Associated p-value** |
| | | Lung vs Breast | 7.36E-04 | 0.003 |
| | | Liver vs Breast | 0.017 | 0.063 |
| | | Lung vs Liver | 0.027 | 0.266 |

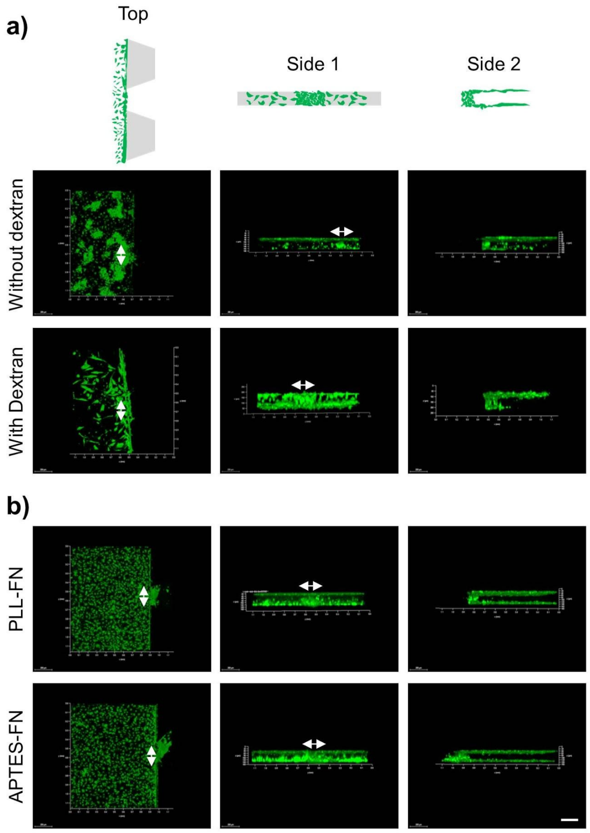


**Figure S1.** APTES and PLL provided a suitable surface for protein coating with homogeneous distribution of endothelial cells in the presence of dextran. a) Schematic representation of top and side views of a post gap on the EX-chip. 3D confocal images showing the distributions of endothelial cells (green) with or without dextran. The endothelial monolayer between the post gaps are represented with two-sided arrows (Scale bar: 200 µm). b) 3D top and side images of EX-chip showing endothelial cells (green) loaded on PLL-FN and APTES-FN coated surfaces. The endothelial monolayer between the post gaps are represented with two-sided arrows (APTES: 3-Aminopropyl triethoxysilane, PLL: Poly-L-lysine solution, FN: fibronectin) (Scale bar: 200 µm).
